# AI predictions and the expansion of scientific frontiers: Evidence from structural biology

**DOI:** 10.64898/2026.04.06.716821

**Authors:** Mengyi Sun, Sukwoong Choi, Yian Yin

## Abstract

Artificial intelligence holds the potential to expand the frontier of scientific research, yet recent work has raised concern that it may instead narrow scientific attention to well-established areas. Here, leveraging the 2021 release of AlphaFold2 as a quasi-experimental opportunity, we provide field-level evidence that AI can redirect collective attention toward more novel research targets. Tracking 245,396 experimental structures in the Protein Data Bank, we show that a long-running decline in the study of novel proteins halted after AlphaFold2’s release, with the shift concentrated among studies citing AlphaFold2 and targets with high-confidence predictions. This pattern extends to 248,191 downstream papers that consume structural knowledge, where engagement with genes lacking experimental structures and with understudied human genes increased since 2021. Amid rising concern that AI may reinforce scientific canons, our findings offer an early field-level case where AI predictions expand scientific frontiers, consistent with the idea that the real-world consequences of AI on science depend on where their informational gains are greatest. These results suggest AI can complement human knowledge and redirect collective attention in science, with broad implications for emerging AI for science models.

---

Science progresses through sustained exploration of the unknown (*1*–*4*). Yet as the burden of knowledge accumulates, the frontier becomes prohibitively costly to expand (*5, 6*). Growing concerns have emerged that scientific attention is increasingly concentrated on familiar questions and well-explored paradigms (*7, 8*), leading to a documented decline in scientific novelty and disruptiveness across disciplines(*9, 10*). This contraction, if continued, may jeopardize the recognition and nurturing of emerging scientists (*11*), undermine the long-term vitality of the R&D ecosystem (*12*), and threaten the pipeline of breakthroughs that fuels economic growth and human prosperity (*13*).

Artificial intelligence offers a potentially powerful, yet contested, mechanism (*14*) to counter this drift. By generating predictions at scale, AI systems may drastically lower the cost of exploring unknown areas and expand the horizon of viable targets. Recent models (*15*–*17*) increasingly learn transferable regularities that can generalize beyond observed samples, raising the prospect that their predictions can prove informative in the broader space and empower researchers to venture beyond the established canon (*18*–*23*). However, recent empirical evidence warns of a darker possibility: while AI can boost individual productivity, it may simultaneously narrow the collective diversity of ideas (*24*–*28*). Driven by institutional incentives, competitive environments, and resource constraints, scientists may strategically converge on data-rich areas where models perform best, further reinforcing existing paradigms (*29, 30*).

Despite these promises and perils, there has been limited large-scale evidence documenting whether and how AI can expand the scientific frontier in the wild. Although AI models have grown increasingly capable, a translation gap from computational predictions into real-world discovery persists, and adoption rates for frontier models—particularly in experimental wet labs—have remained too low to demonstrate macro-level shifts. We address this empirical challenge by leveraging the 2021 release of AlphaFold2 (*16, 31*) (AF2) as a unique quasi-experimental design. As one of the first widely adopted AI-for-science tools, AF2 provides structure predictions for over 200 million proteins, bringing predictive power at unprecedented scale into a field historically dominated by the intensive exploitation of familiar targets and a vast expanse of structural dark matter. Moving beyond recent studies of individual adopters (*32*–*34*), we examine whether AF2 altered research directions in structural biology and beyond.

## Results

### AI adoption in structural determination

How rapidly has AI been adopted in determining protein structures? We collected and analyzed large-scale data on all experimental structures deposited in the Protein Data Bank (*35*), including 245,396 PDB entries and 87,656 associated papers (SM S1.1). We further use citations to key AlphaFold papers as a proxy for AI use in experimental structure determination (SM S1.4, S2.1). Figure 1A, B document rapid uptake shortly after the tool became available. Although only four years have passed since AF2’s debut, 20.5% of new structures deposited in 2024 are now associated with papers citing AlphaFold2. Adoption is markedly higher among cryo-electron microscopy (cryo-EM) entries (38.7%), nearly doubling the rate for X-ray crystallography (20.2%). Notably, in both modalities, adoption follows highly nonlinear dynamics: initial acceleration followed by slower growth and eventual saturation. The S-shaped curve is well captured by a logistic diffusion model; extrapolating the curve from empirical data up to late 2022 (18 months post-AF2) is sufficient to predict subsequent saturation (SM S2.1). Despite the unprecedented speed of AlphaFold’s uptake, its longer-term diffusion appears to follow the same trajectory as many earlier technological tools (*36*).

**Fig. 1.**
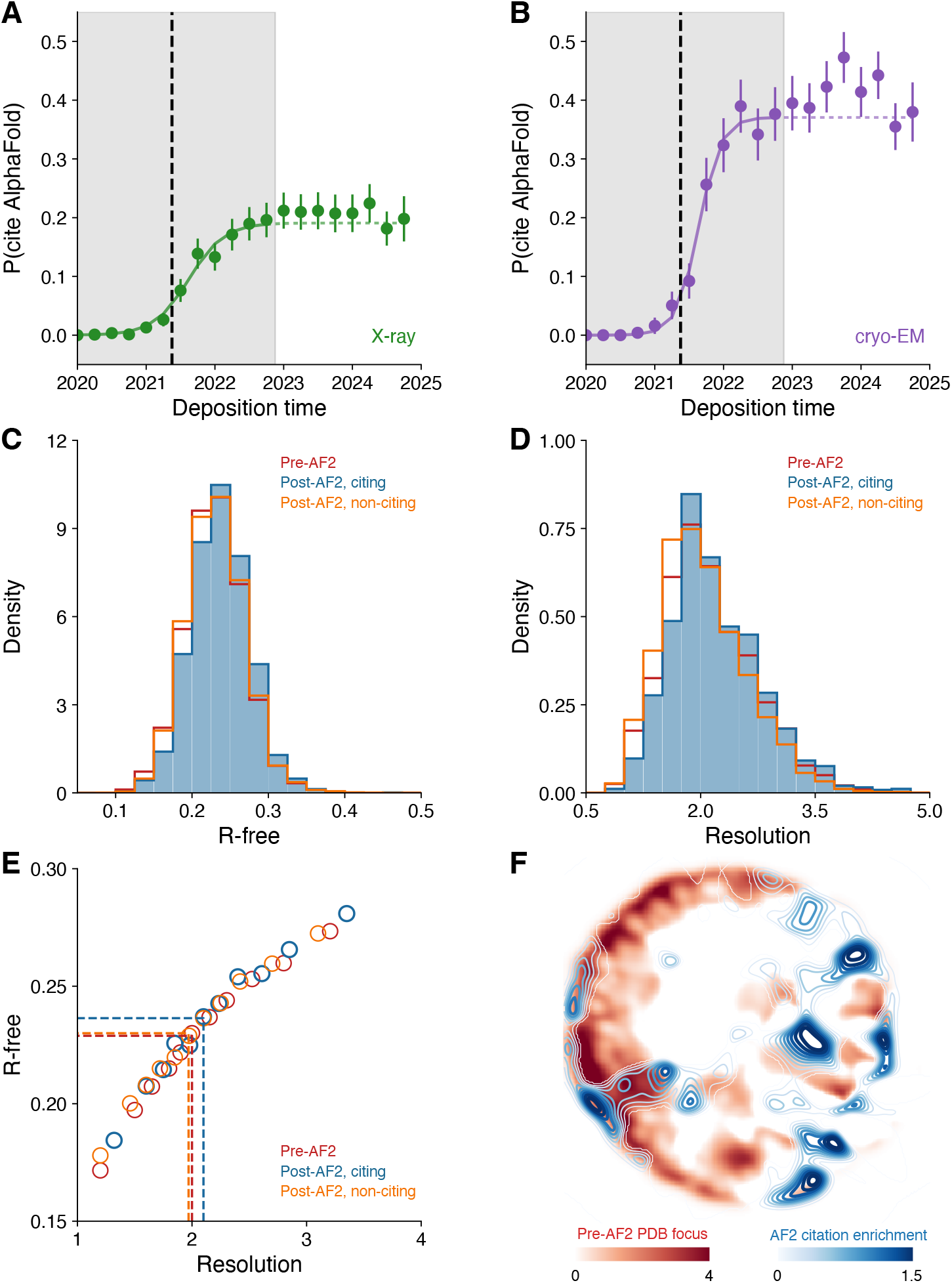
Adoption of AlphaFold2 in structural biology. **A–B**, Fraction of new PDB structures associated with papers citing AlphaFold2, shown separately for X-ray crystallography (**A**) and cryo-electron microscopy (**B**) by deposition quarter. Points show empirical data; curves show fitted logistic diffusion models estimated on data through late 2022. **C**, Distribution of R-free values for structures deposited before AF2 release (pre-AF2; red), and post-AF2 structures in papers citing AlphaFold2 (post-AF2, citing; blue) or not citing AlphaFold2 (post-AF2, non-citing; orange). **D**, Distribution of crystallographic resolution across the same three groups. **E**, Mean R-free as a function of resolution, binned by resolution quantile. Dashed lines indicate group medians. When conditioned on resolution, the three groups collapse onto a single curve, indicating that the apparent quality deficit in AF2-citing structures is fully explained by a compositional shift toward harder targets. **F**, Two-dimensional projection of protein sequence embedding space (ProtTrans T5 XL U50), with PaCMAP applied to PCA-reduced AlphaFold embeddings as the reference manifold. Red shading indicates density enrichment of pre-AF2 PDB entries relative to the AlphaFold universe; blue shading indicates density enrichment of post-AF2 AF2-citing PDB entries relative to post-AF2 non-citing entries. AF2-citing entries occupy regions of sequence space largely distinct from the pre-AF2 structural canon. Error bars in all panels represent 95% confidence intervals.

The rapid adoption prompts us to ask: does the growing volume of AI-assisted research come at a cost to experimental quality? The quality effect of AI for science remains debated in the literature (*37*–*39*), largely because of the challenges in measuring scientific quality at scale. Here we leverage a unique quality measure for PDB structures: R-free (*40*), which quantifies the agreement between the diffraction data and the refined model. Fig. 1C plots the distribution of R-free for structures citing AI algorithms alongside contemporaneous non-citing entries and pre-AF2 entries. AF2-citing entries exhibit higher (worse) R-free values, suggesting an apparent net loss in average quality.

Why are AI-powered discoveries associated with lower quality? We further explore this apparent puzzle by examining a second quality indicator: resolution, which measures the precision of initial diffraction data. Resolution is determined primarily by crystallization and imaging, and thus unlikely to be substantially affected by predictive algorithms. Figure 1D examines the distribution of resolution across the three groups and shows that AF2-using structures also link to lower-quality initial data (*P* < 0.001). More importantly, when we condition on resolution, all three groups collapse onto a single curve (Fig. 1E). Controlling for initial data quality (resolution), AlphaFold use is associated with neither gains nor losses in outcome quality (*P* = 0.980). The apparent quality deficit therefore suggests a compositional shift: AI use is not uniform in the protein space. Rather, it appears preferentially deployed on challenging cases with inferior diffraction data, which is sufficient to account for the R-free difference. These results highlight that AI use is more closely tied to changes in research focus than to quality. A sequence embedding-based visualization of the protein space is consistent with this interpretation (Fig. 1F, SM S2.2,2.3): AlphaFold-citing entries enrich regions of sequence space largely distinct from those densely populated by pre-AF2 PDB entries, indicating that AF2 redirected attention toward proteins beyond the established structural canon.

### Directional shifts in experimental structures

To systematically measure changes in research focus, we construct entity-level measures of novelty by grouping PDB entities into sequence-similarity clusters (70% identity, alignment-based) (*41, 42*) (SM S2.4). We then classify a PDB structure deposited in year *t* as novel if no prior entities targeted the same cluster before *t* – Δ*t*, where Δ*t* = 1 year to account for the lag between project initiation and deposition (SM S2.6). The relationship between research projects and PDB entities is often not one-to-one, as a single project may yield multiple PDB entities. To this end, we group all entities deposited by the same team on the same day into a single project, assigning equal fractional weights within each project (SM S2.7, Fig. S1).

Figure S2 plots novelty, defined as the average fraction of novel structures targeted per project, over time. Consistent with existing analysis (*43*), we find a historical decline through the early 2020s. Yet in more recent years, the decline halted, where the inflection appears to align well with the public release of AF2 and the availability of its large-scale prediction. To test this relationship more rigorously, we estimate a segmented linear regression examining changes in slope around AF2’s release (Fig. 2A, SM S2.10). The results suggest a significant slope change: prior to AlphaFold the novelty rate declines by 1.2 percentage points per year (*P* < 0.001), a trend that halts and becomes statistically indistinguishable from zero (*P* = 0.590) in the post-AF2 period (Fig. 2A).

**Fig. 2.**
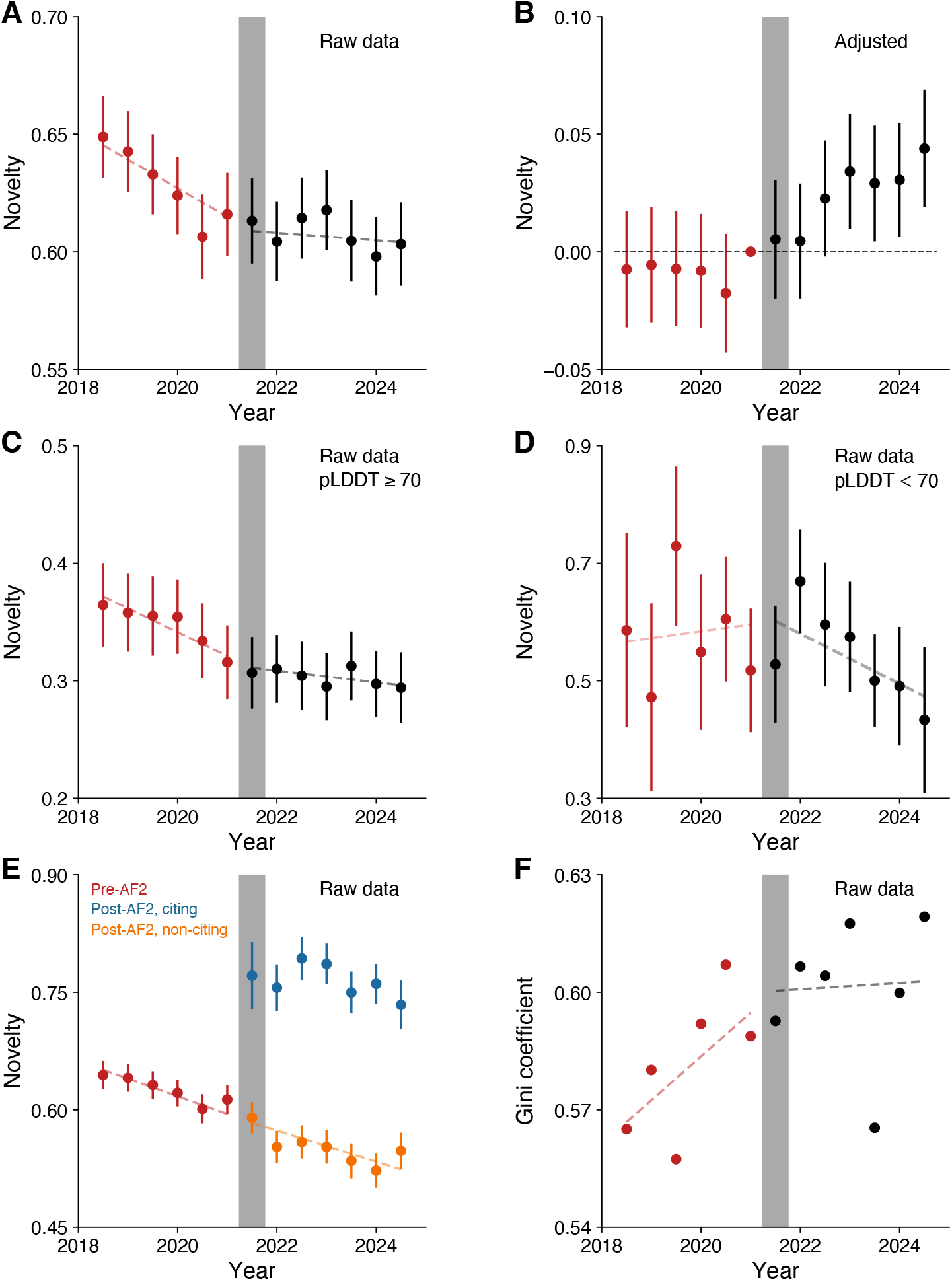
AlphaFold2 arrests a long-running decline in the novelty of experimental structural targets. **A**, Fraction of novel protein clusters targeted per PDB project, by deposition time (6-month bin). A project is classified as targeting a novel cluster if no prior PDB entry mapped to the same 70% sequence-identity cluster before the preceding year. Red points indicate pre-AF2 period; black points indicate post-AF2 period. Dashed lines show segmented linear regression fits. Prior to AF2, novelty declined by 1.2 percentage points per year (*P* < 0.001); post-AF2, the slope is statistically indistinguishable from zero (*P* = 0.590). **B**, Event-study representation of novelty deviations relative to the extrapolated pre-AF2 linear trend. **C–D**, As in (**A**), stratified by protein cluster with AlphaFold2 prediction pLDDT ≥ 70 (**C**) or pLDDT < 70 (**D**). The novelty inflection appears concentrated in the high-confidence regime, yet absent in the low-confidence regime. **E**, As in (**A**), stratified by whether the depositing paper cites AlphaFold2. AF2-citing papers target 38.3% more novel sequences than non-citing contemporaneous papers; no significant inflection is observed among non-citing papers. **F**, Gini coefficient of PDB research intensity across 70% sequence-identity clusters, calculated in 6-month bins. The Gini coefficient grew significantly in the pre-AF2 period (*P* = 0.009) but has been statistically stable thereafter (*P* = 0.718), indicating an arrest of increasing concentration. Error bars in all panels represent 95% confidence intervals.

Further, using a linear approximation of the pre-existing trend (Fig. 2B), we estimate that by late 2024, the rate of novel PDB structures is 4.4 percentage points higher than expected. This link between AI use and novel research is striking, as most AI algorithms are trained on existing data derived from prior research and often struggle where training examples are sparse. If scientists leveraging AI predictions were to prioritize targets with accurate predictions, one might expect research to increasingly converge to well-explored areas (*7, 28, 44, 45*). Yet we observe the opposite pattern: AI predictions have enabled exploration into previously uncharacterized regions of the protein space, directing experimental effort toward proteins that lacked prior structural data.

Several lines of further analyses support this interpretation. First, the effect is more pronounced when the algorithm yields high-quality predictions. To this end, we exploit a key feature of AF2: the predicted Local Distance Difference Test (pLDDT) score (*16, 46*), a residue-level confidence estimate that is intrinsic to the prediction. On a 0–100 scale, scores above 70 are often considered comparable to experimental accuracy. Indeed, for clusters that can be directly mapped to AlphaFold predictions, the arrest of novelty decline is concentrated in those with high-pLDDT predictions (Fig. 2C) and largely absent in those with low-pLDDT predictions (Fig. 2D).

Further, the elevated novelty is predominantly contributed by entries whose corresponding papers explicitly cite AlphaFold (Fig. 2E). PDB papers citing AlphaFold engage 38.3% more novel protein sequences than non-citing papers from the same period, again pointing to a close link between AI use and expansion into unexplored sequence space. Yet comparing non-citing papers before and after AF2 release, we find no statistical evidence for the nonlinear transition (*P* = 0.639), suggesting that the macro-level shift cannot be simply explained by alternative temporal factors.

Overall, the increase in research novelty is not spread evenly across the structural dark matter, but concentrated when scientists actively deploy AI tools and where AI provides confident predictions. Amid ongoing discussion about whether predictions might substitute for experiments, our results suggest the opposite: structural biologists selectively use AlphaFold predictions to pursue novel proteins precisely where the AI is confident, indicating a complementary rather than substitutive relationship (*42, 47*). SM S4 documents further robustness across different analytical choices and thresholds, showing our results remain remarkably consistent (Fig. S4-S7).

The novelty change documented in Fig. 2A-E also implies a broader redistribution of structural research effort: as researchers increasingly examine proteins beyond existing literature, the research ecosystem may see an increase in target diversity. To this end, we calculate, for each 6-month time period, the Gini coefficient of PDB research intensity (SM S2.9). Consistent with this hypothesis (Fig. 2F), while the Gini coefficient had grown over time (*P* = 0.009), it has largely stalled post-AF2 (*P* = 0.718).

Together, these results point to a directional shift that goes beyond the productivity and quality effects of AI: as AF2 converts a substantial fraction of unknowns into knowns, scientists start redirecting attention toward a diverse set of promising directions.

### The directional shift propagates to downstream research

The directional shift among experimental producers raises a further question: can AF2 further impact the broader community that consumes structural knowledge in downstream research? Protein structures inform a wide range of biomedical tasks, such as drug design (*48*), functional annotation (*49*), and mechanistic modeling (*50*). Thus, the population of experimental *producers* examined above represents only one side of the ecosystem, and reallocation of structural effort toward novel targets may have limited impact if downstream *consumers* continued to draw selectively on well-characterized proteins.

To answer this question, we used NCBI PubTator3 (*51*), a text-mining system that maps biomedical concepts to entity identifiers in PubMed titles and abstracts, linking 2.89M papers with 132K genes from 2015 to 2025 (SM S1.3). For each paper, we used GPT-4o-mini to classify whether it likely consumed structural information (see SM S2.5 and Fig. S3 for classification details and validations against a hand-labeled subset). Figure 3A plots the fraction of papers classified as consuming structural knowledge, which follows a modest increasing trend before AF2’s release (0.10 percentage points per year) but accelerates markedly thereafter (0.46 percentage points per year). Tracing citations within these papers, the disproportionate post-2021 increase is largely associated with papers citing AlphaFold, suggesting an extensive margin expansion in AF2-related downstream research (Fig. S8A).

**Fig. 3.**
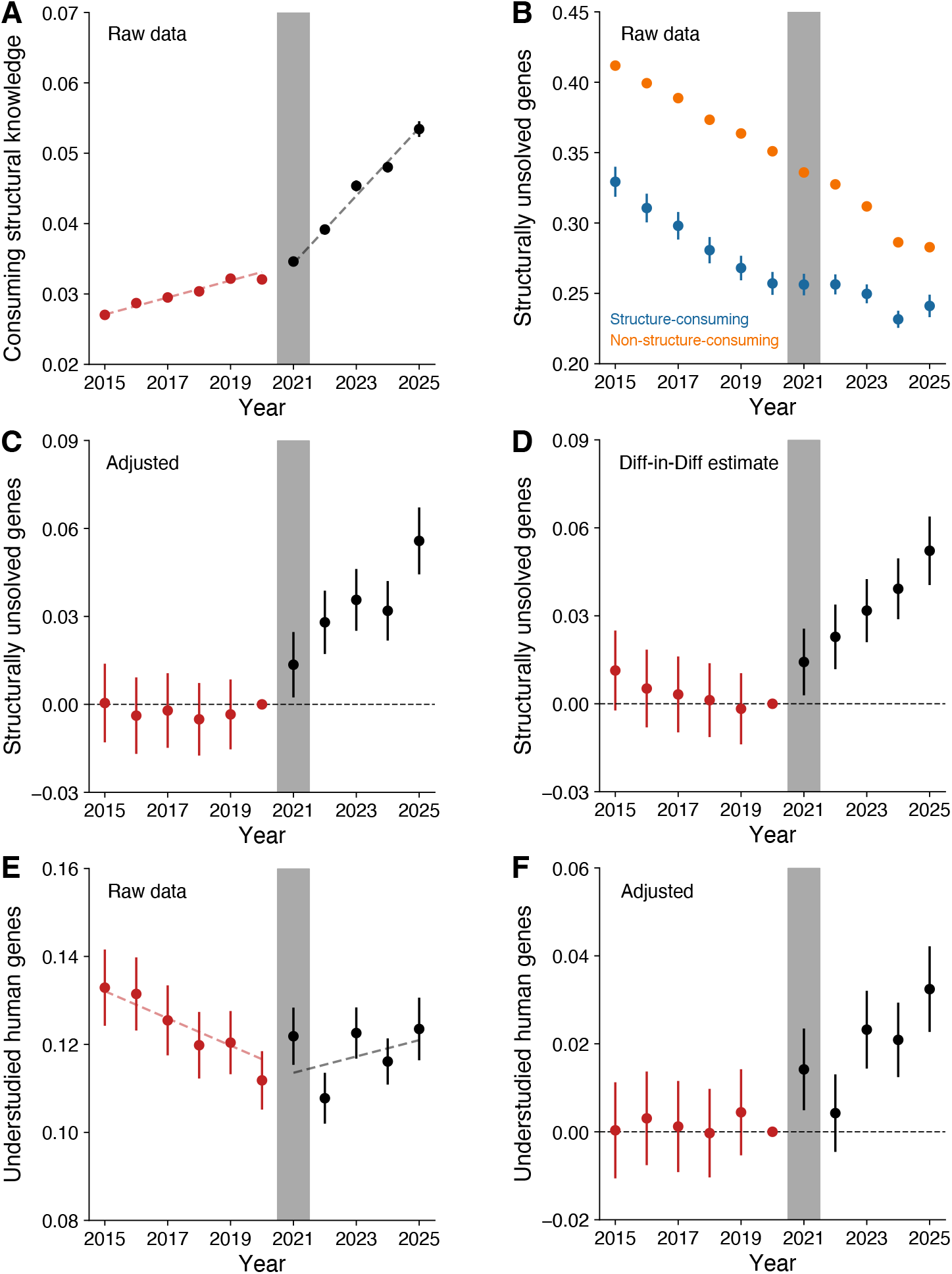
The directional shift propagates to broader biomedical papers consuming structural knowledge. **A**, Fraction of PubTator3-annotated gene-mentioning papers classified as structural knowledge consuming by publication year, following a marked acceleration post-AF2 (0.46 percentage points per year). **B**, Among structure-consuming (blue) and non-consuming (orange) papers, the fraction engaging genes without any experimentally released structure at the time of publication. Structural consumers show a sustained pre-2021 decline that reverses after AF2’s release, while non-consumers continue to decline. **C**, Event-study estimates of deviation from pre-AF2 trend in the fraction of unsolved genes targeted by structure-consuming papers. By 2025, the rate is 5.58 percentage points above the pre-trend extrapolation. **D**, Difference-in-differences estimates using non-structure-consuming gene-mentioning papers as controls. The two groups exhibit parallel pre-trends through 2020; structure-consuming papers diverge significantly thereafter (5.22 percentage points higher by 2025). **E**, Fraction of structure-consuming papers engaging genes identified as potentially understudied in the human genome. **F**, Event-study estimates of deviation from pre-AF2 trend in understudied gene engagement, corresponding to a 3.24 percentage point increase. Error bars in all panels represent 95% confidence intervals.

On the intensive margin, we further examine which genes scientists engage with. Paralleling the novelty definition in Fig. 2, we focus on an extreme case: genes without any experimental structure released by the year of the consumer paper’s publication (SM S2.6,2.8). At the individual level, papers citing AF2 study proteins without prior experimental structures at 3.11 times the rate of non-citing papers (Fig. S8B). Consistent with this substantial contrast, Fig. 3B shows an observable system-level shift: within structure-consuming papers, the engagement of structurally unsolved genes follows a persistent pre-2021 decline that has been interrupted sharply thereafter. An event study adjusting for pre-trend (Fig. 3C) estimates an increase of 5.58 percentage points by 2025, representing 21.7% relative growth over the pre-AF2 baseline. To examine whether this can be explained by alternative temporal confounds unrelated to structural knowledge, we further implement a difference-in-differences design, using gene-mentioning papers that do not consume structural information as controls (Fig. 3D). The two groups exhibit parallel trends through 2020, after which structural consumers start targeting structurally unsolved genes at a significantly higher rate, with a magnitude comparable to the event study estimate (5.22 percentage points by 2025).

The growing focus on structurally unsolved genes has important implications along biologically meaningful dimensions. As an illustrative example, we examine whether AF2 redirected attention toward valuable but historically underserved targets in the human genome. Drawing on a machine learning model that predicts expected research intensity from gene-specific chemical, physical, and biological features (*52*), 6,981 out of 15,056 human genes have been identified as potentially understudied in the early 2010s. In our data, the fraction of consumer papers engaging these genes continued to decline until 2021 but stabilized thereafter (Fig. 3E). Compared with the 2020 baseline, we estimate a 29.0% relative increase in research on understudied genes (3.24 percentage points, Fig. 3F), which suggests a substantial diversification effect within the human genome. Together, these findings provide initial evidence consistent with AF2’s directional impact across the broader biomedical literature, highlighting an expansion in the volume of structural engagement and the scope of research targets.

### A minimal model of AI predictions and research directions

Our results may seem paradoxical. AI models are trained on existing data, and a large body of recent work warns that such systems tend to reinforce attention on well-studied targets, concentrating effort where training examples are densest rather than expanding into the unknown (*18, 28, 53*). Why, then, does AF2 produce the opposite pattern? The answer, we argue, lies not in whether a tool uses AI, but in where it delivers its greatest informational contribution.

To formalize this idea, we develop a simple framework (Figure S9A) in which a researcher chooses among projects (e.g., protein sequences) with varying levels of distance from the existing knowledge space, denoted by *z*, where higher *z* corresponds to thinner prior evidence and greater baseline ambiguity (e.g., fewer close homologs). Each researcher selects a single project (or target), maximizing the expected payoff from discovery—the probability of success times the value upon discovery—minus the cost of experimental pursuit. Because only one project can be pursued, choosing a more attractive project raises the opportunity cost of all others. The model suggests that the impact of AI predictions on research directions depends not only on their own predictive power, but also on how that power is distributed across the knowledge landscape. We define the tool’s information gain as *r*(*z*) = *h*^*post*^(*z*) / *h*^*pre*^(*z*), the ratio of (perceived) success probabilities with and without AI. When *r*(*z*) declines with *z*, gains concentrate in data-rich regions and raise the opportunity cost of frontier work—a pattern we term *data-reinforcing*. When *r*(*z*) rises with *z*, gains accrue disproportionately to hard, data-sparse targets, reducing the risk premium on novel work and shifting effort outward—a pattern we term *frontier-expanding*.

This framework shifts attention from whether a tool uses AI to where its informational lift is concentrated across the knowledge landscape. AF2 appears broadly consistent with the frontier-expanding pattern: its architecture appears to capture transferable physical and geometric constraints that generalize beyond close homologs, delivering greater relative gains where prior evidence is thinnest (Fig. S9B). An interesting contrasting case is molecular replacement (MR) (*54*), a widely used technology that uses known structures as templates to solve the crystallographic phase problem (*29*). MR is particularly powerful for proteins with close homologs but adds little value at higher *z*, which appears consistent with the data-reinforcing pattern (Fig. S9C). Full model derivation and connections to empirical proxies are documented in Supplementary Material S3.

## Discussion

Our findings provide field-level evidence that AI predictions can redirect scientific effort toward new frontiers. In structural biology, a persistent decline in the targeting of novel proteins halted after the release of AF2. The shift is concentrated where AF2 should matter most—among studies that use it, for targets with high-confidence predictions, and in downstream research that consumes structural knowledge. While AI is often treated as a productivity tool that accelerates existing workflows, our results suggest a broader function: it can reorganize which problems a field regards as tractable and worth pursuing.

This paper complements a growing literature that has mainly examined how AI affects individual scientists or labs, comparing research behaviors between adopters and non-adopters ^32,33,55–57^. By treating the field as the unit of analysis, we identify a collective effect: once predictive tools diffuse broadly enough, they can reshape the distribution of attention across a field. AF2 offers a unique empirical opportunity, combining a quasi-experimental design, rapid diffusion, and observable downstream impact. But the same patterns will likely emerge as AI models across domains become increasingly integrated into scientific workflows and capable of learning and generalizing transferable scientific regularities.

The directional change documented here has broad implications for how effects of AI in science should be evaluated. AF2-linked structures appear to underperform on experimental quality measures, which reflects a compositional shift in the problems being attempted: scientists use AI on problems that appear intrinsically harder. More generally, when a technology induces selection into more challenging problems, standard metrics such as quantity and quality can mislead if interpreted without regard to that selection. The same logic applies to novelty. In a setting where novelty has been declining, arresting that decline is itself a meaningful shift, and analyses that focus only on the post-period levels may risk understating the true impact ^58^.

These findings further suggest AI need not simply reinforce attention on scientific canons, even when trained on data generated by past research. Our minimal modeling framework is useful in integrating these seemingly contradictory perspectives (Fig. S9): for scientific progress, it may matter at least as much where a tool’s informational gain is concentrated as how well it performs on average. An AI system can score highly on benchmarks while concentrating its gains in already-tractable regions, thereby reinforcing reliance on well-established areas. Conversely, a tool with modest average performance can be consequential if it extends useful guidance into data-sparse domains. In our case, AF2 appears more consistent with a frontier-expanding pattern (Fig. S9B), whereas molecular replacement more closely reflects a data-reinforcing pattern (Fig. S9C, also see Ref (*29*)).

There are several limitations that readers should keep in mind. We do not claim a perfectly isolated causal estimate, as AF2 emerged alongside other changes in biomedical research. While we have excluded COVID-related research ^59^ in our analysis, we cannot fully rule out the role of the post-COVID resumption of laboratory research. Further, other relevant AI technologies, including protein language models ^60,61^, also emerged in the same period. However, the null effect among low-pLDDT targets, the high novelty of AF2-citing studies, and the divergence between papers that do and do not consume structural knowledge, appear difficult to reconcile with a simple background temporal trend.

Even with these caveats, the broader lesson remains: beyond boosting quantity and quality, AI can matter for scientific progress through where it makes exploration more feasible. With human scientists in the loop, AI predictions need not fully replace experiment to be important. Rather, it may be sufficient that they complement human efforts ^62^ by providing guidance where prior knowledge is sparse and the perceived risks of exploration are high. AF2 showcases an early and clear view of this broader dynamic: it helped arrest a long-running novelty decline, even without matching experimental standards across all targets ^63,64^. As foundation models and AI agents proliferate ^65^ across mathematical ^66,67^, physical ^68^, and life sciences ^69^, their long-run impact on science may depend not only on how accurately they predict, but also on how far beyond existing concentrations of knowledge they remain useful.

## Acknowledgments

We thank Juan Mateos Garcia, George Richardson, Ryan Hill, and Carolyn Stein, as well as seminar participants at Google DeepMind, Cold Spring Harbor Laboratory, and the ICML AI4Science workshop for helpful correspondence and discussions.

## Author contributions

Conceptualization: MS, SC, YY Methodology: MS, SC,YY Investigation: MS, SC, YY Visualization: MS, YY Funding acquisition: YY Project administration: YY Supervision: YY Writing – original draft: MS, SC, YY Writing – review & editing: MS, SC, YY

## Data, code, and materials availability

Data and code for reproduce the analysis would be made available upon publication.

## Supplementary Materials

Materials and Methods

Supplementary Text

Figs. S1 to S9

Tables S1 to S2

References (*1*–*20*)

